# Striatal neuron excitability is regulated by huntingtin in the adult brain

**DOI:** 10.1101/2025.08.14.670392

**Authors:** Jessica C. Barron, Meghan L. Greenland, Fatemeh Ashrafganjoie, Emily P Hurley, Firoozeh Nafar, Craig S. Moore, Matthew P. Parsons

## Abstract

Huntington’s disease (HD) is a hereditary neurodegenerative disease that typically presents during midlife and is characterized by a combination of motor, cognitive and psychiatric symptoms. HD is fatal and arises from a mutation in the huntingtin (*HTT*) gene, which results in decreased neuronal health followed by brain atrophy, with spiny projection neurons (SPNs) of the striatum being especially vulnerable to degeneration. HTT loss-of-function, caused by haploinsufficiency of the wild type HTT gene (wt*HTT*), is an important feature of HD pathophysiology that has previously been understudied compared to mutant HTT gain-of-function mechanisms. wtHTT is essential for nervous system development and functions as a scaffolding protein to support many vital cellular functions including axonal transport, autophagy and synaptic plasticity. Here, we examined the consequences of wtHTT deletion in the adult striatum by conditionally inactivating wtHTT in 2-4 month old male and female *Htt*^fl/fl^ mice. wtHTT loss of function in mature SPNs decreased intrinsic neuronal excitability and produced a neuroinflammatory response in these mice, while tissue organization, spine morphology and motor behaviour remained unaffected. Results presented here provide additional evidence that wtHTT is vital for maintaining neuronal health in the adult brain and highlight some potential adverse consequences of non-selective HTT-lowering for the treatment of HD.

## Introduction

Huntington’s disease (HD) is a monogenic neurodegenerative disease caused by an expansion of 35 or more CAG repeats in the huntingtin (*HTT*) gene, which leads to the production of a pathogenic mutant HTT protein, and has an estimated prevalence of 13.7 per 100,000 in the general population (Fisher and Hayden, 2014). Individuals with HD experience a triad of motor, cognitive and behavioural symptoms, which typically present during middle age; however, symptom onset can vary from childhood to late adulthood (Walker, 2007). Despite many ongoing clinical trials, there is currently no cure for this fatal disease (Tabrizi et al., 2022). HD results in widespread atrophy throughout the brain, with spiny projection neurons (SPNs) of the striatum being particularly vulnerable (Vonsattel et al., 1985; G. Vonsattel and DiFiglia, 1998). In line with human pathology, HD mouse models recapitulate SPN-specific neurodegeneration (Hodgson et al., 1999; Slow, 2003; Heikkinen et al., 2012). Whole-cell recordings from HD SPNs display alterations in excitatory activity early in the disease course (Li et al., 2004; Milnerwood et al., 2010, 2012). By DIV 21, SPNs co-cultured with cortical neurons from the YAC128 HD mouse model have reduced dendritic arborization, fewer excitatory events and fewer readily releasable pool synaptic vesicles (Buren et al., 2016). *Ex vivo*, many HD mouse models display similar alterations in SPN electrophysiological properties, such as fewer sEPSC and greater spontaneous IPSC events (Cummings et al., 2010). Comparable results are also seen in whole-cell recordings from striatal slices of Q175 knock-in HD mice (Indersmitten et al., 2015), highlighting these electrophysiological changes as a feature of HD brain pathology.

While mutation of the HTT protein has profound gain-of-function pathogenic effects on striatal neurons, loss-of-function of wild type HTT (wtHTT) has also been shown to influence SPN health. For instance, striatal neurons do not produce the essential neurotrophin, BDNF, and therefore rely on its transport from the cortical projection neurons (Altar et al., 1997). wtHTT loss of function has been shown to downregulate transcription of BDNF as well as disrupt its anterograde and retrograde axonal transport (Zuccato et al., 2001, 2003; Gauthier et al., 2004; Colin et al., 2008; Her and Goldstein, 2008). Non-pathogenic HTT is also known to have an antiapoptotic effect in HD culture and mouse models (Leavitt et al., 2001, 2006). Further, previous research has shown that deletion of wtHTT in YAC128 mice worsened striatal atrophy, exacerbated motor deficits and reduced survival rates (Van Raamsdonk et al., 2005a). Although overexpression of wtHTT reduced striatal atrophy, this did not translate to a complete rescue of motor deficits in YAC128 animals (Van Raamsdonk et al., 2006). We have previously reviewed the extensive literature that supports a major role for wtHTT at the synapse (Barron et al., 2021). Notably, deletion of wtHTT disrupts corticostriatal excitatory synapse development where wtHTT-depleted neuronal networks were initially hyperexcitable at postnatal day 21, followed by a reduction in excitatory synapse activity and density, spine maturity and dendritic complexity by 5 weeks of age (McKinstry et al., 2014). wtHTT is also a critical regulator of presynaptic neurotransmission in the striatum, as its deletion in striatal cultures disrupts synaptic vesicle endocytosis (McAdam et al., 2020). Other groups have investigated the embryonic deletion of wtHTT in distinct subpopulations of striatal neurons, such as in D1 and D2-expressing SPNs, and found it negatively impacted DARPP-32 levels and motor behaviour (Mehler et al., 2019; Burrus et al., 2020). These studies provide compelling evidence for wtHTT as a necessary regulator of SPN survival in the HD brain.

Conditional deletion of wtHTT in the dorsal hippocampus of 2-4 month *Htt*^fl/fl^ mice results in profound morphological deficits and widespread reactive gliosis within two months (Barron et al., 2025). wtHTT-depleted CA1 pyramidal neurons also had reduced intrinsic excitability, increased spontaneous EPSCs and loss of NMDAR-dependent long term potentiation (LTP) accompanied by impaired spatial memory (Barron et al., 2025). These findings uncovered a novel role for wtHTT as an essential regulator of hippocampal synaptic plasticity and survival. As the striatum is a particularly susceptible brain area in HD and striatal neurons are known to be negatively impacted when wtHTT is deleted embryonically, we hypothesized that wtHTT deletion in adult striatal SPNs would negatively impact neuronal function and lead to neuroinflammation. Here, the Cre-lox system was used to inactivate *HTT* in striatal neurons of 2-4 month old male and female *Htt*^fl/fl^ mice. wtHTT conditional knockout (cKO) lead to significant astrogliosis and decreased intrinsic neuronal excitability in these mice, implicating wtHTT as an important regulator of SPN health in the adult mammalian brain.

## Materials & Methods

### Animals

wtHTT cKO mice were initially provided by Dr. Scott Zeitlin, University of Virginia, Charlottesville, VA (Dragatsis et al., 2000). Breeding colonies were established and maintained at [Author University] Animal Care facility. Mice used in these experiments have their endogenous *wtHtt* alleles flanked with loxP sites to conditionally inactivate the mouse *Htt* gene in a homozygous manner by Cre expression (*Htt*^fl/fl^). Mice were bred as homozygous x homozygous. Equal numbers of male and female mice were used for all experiments. Mice were group housed in ventilated cage racks and kept on a 12 h light/dark cycle (lights on at 7:00 A.M.) with food and water available *ad libitum*. All animal procedures were performed in accordance with the [Author University] animal care committee’s regulations.

### Western blotting

For Western blot experiments, the dorsal striatum of AAV-Cre-eGFP or AAV-eGFP-injected *Htt*^fl/fl^ mice were dissected and homogenized in 400 µl of lysis buffer containing halt protease and phosphatase inhibitor cocktails (Thermofisher Scientific, catalog #78440). The supernatant was collected, and the protein concentration was determined using bicinchoninic acid standards (Pierce Bicinchoninic Acid Protein Assay Kit, Thermofisher Scientific, catalog #23227). Protein samples were prepared by adding 4× lithium dodecyl sulfate sample buffer (Invitrogen, catalog # B0007) and heating at 70°C for 10 minutes. Samples were loaded onto a 4–12% Bis-Tris Plus gel (Invitrogen, catalog #NW04122BOX) for electrophoresis. Following electrophoresis, proteins were transferred onto a 0.45 um nitrocellulose membrane (Santa Cruz Biotechnology, catalog #sc-201706) using wet transfer at 24 V for 16 hours at 4°C. Transfer buffer contained 10% methanol. After transfer, the membrane was blocked with 5% non-fat milk in Tris-buffered saline with 0.05% Tween-20 for 1 hour at room temperature to prevent nonspecific binding. Primary antibodies for anti-HTT (1:1000; mouse, Sigma Aldrich, catalog #MAB2166) and anti-beta-actin (1:5000, mouse, Sigma Aldrich, catalog #A5316 clone AC74), and a goat anti-mouse IgG secondary antibody (1:5000, Thermofisher Scientific, catalog #31430) were used. Blots were developed using a chemiluminescent horseradish peroxidase substrate (Super Signal West Pico Plus, Thermofisher Scientific, catalog #34580). Band densities were quantified in FIJI (Schindelin et al., 2012), and HTT band densities were normalized to actin.

### Stereotaxic surgery

*Htt*^fl/fl^ mice aged 2-4 months were anesthetized using isoflurane (3% induction, 1.5–2% maintenance) and received a single subcutaneous injection of 2 mg/kg meloxicam in the abdomen and a single 0.1 ml/0.2% lidocaine injection below the scalp. Two small holes were drilled in the skull at the target brain coordinates using an Ideal Micro-Drill (Harvard Apparatus). A Model 7002 KH Neuros Hamilton Syringe and an infusion pump (Pump 11 Elite Nanomite, Harvard Apparatus) were used to inject 1 μl pENN.AAV.hSyn.HI.eGFP-Cre.WPRE.SV40 (Addgene, catalog #105540-AAV1) or 1 μl pAAV-hSyn-EGFP (Addgene, catalog #50465-AAV1) bilaterally into the dorsal striatum (injection rate, 2 nl/s). The following coordinates were used with respect to distance from bregma: 0.6 mm anterior, 1.8 mm medial/lateral and 2.7 mm ventral to the brain surface. The syringe was left in place for at least 5 min before slow removal from the brain. Mice received a single 0.5 ml of 0.9% saline injection subcutaneously in the abdominal region, and the scalp incision was sutured. Mice recovered for 20-30 minutes post-surgery on a heating pad before being returned to their home cages. All experiments were performed 1-2 months following stereotaxic surgery.

### Acute slice preparation

Mice were anesthetized using isoflurane before decapitation and whole-brain removal. Brains were immersed in ice-cold oxygenated (95% O_2_/5% CO_2_) NMDG-HEPES artificial CSF slicing solution consisting of (in mM): 92 NMDG, 2.5 KCl, 1.25 NaH_2_PO_4_, 30 NaHCO_3_, 20 HEPES, 25 glucose, 2 thiourea, 5 Na-ascorbate, 3 Na-pyruvate, 0.5 CaCl_2_·2H_2_O, and 10 MgSO_4_·7H_2_O. NMDG solution was adjusted with HCl to reach a pH of 7.3–7.4. Dorsal striatal acute slices from one hemisphere (the opposite hemisphere was used for Golgi staining described below) were sectioned using a Precisionary compresstome. Slices were then transferred into warmed (32–34 °C) NMDG-HEPES artificial CSF and underwent a sodium spike-in protocol (Ting et al., 2018). After this, slices recovered for 30-45 minutes in HEPES holding solution consisting of (in mM): 92 NaCl, 2.5 KCl, 1.25 NaH_2_PO_4_, 30 NaHCO_3_, 20 HEPES, 25 glucose, 2 thiourea, 5 Na-ascorbate, 3 Na-pyruvate, 2 CaCl_2_·2H_2_O, and 2 MgSO_4_·7H_2_O. HEPES artificial CSF solution was adjusted with NaOH to reach a pH of 7.3–7.4. Artificial CSF used during electrophysiological recordings consisted of (in mM) 125 NaCl, 2.5 KCI, 25 NaHCO_3_, 1.25 NaH_2_PO_4_, 1 MgCl_2_, 2 CaCl_2_, 10 glucose, and osmolarity adjusted to be approximately 310 mOsm.

### Whole-cell patch clamp electrophysiology

Whole-cell patch clamp recordings of green fluorescent protein (GFP)-positive striatal neurons were performed using 3-7 MΩ patch pipettes filled with potassium-gluconate internal recording solution made up of (in mM) 123 potassium-gluconate, 2 MgCl_2_, 1 KCl, 0.2 EGTA, 10 HEPES, 5 Na_2_ATP, 0.3 Na_2_GTP, with osmolarity adjusted to 285 mOsm and containing Alexa Fluor 594 Hydrazide (ThermoFisher, catalog #A10442) to visualize successfully patched neurons. Only GFP-expressing neurons were patched. Input-output curves were obtained in current clamp mode, injecting current from -250 pA to + 400 pA in 50 pA increments. Next, cells were held in voltage clamp (Vhold = -70 mV), and two minutes of stable recording was obtained to analyze sEPSC frequency and amplitude. Whole-cell recordings were analyzed using ClampFit version 10.6. Detected sEPSCs with amplitudes less than 5 pA were removed from the data analysis.

### Golgi staining

Golgi staining was performed using Bioenno Lifesciences *sliceGolgi Kit* (catalog #003760) materials and protocol. Mouse hemibrains dissected during striatal acute slice preparation described above and were then submerged in a fixative buffer containing formaldehyde and glutaraldehyde. Brains remained in fixative solution at 4 ℃ overnight. Fixed brain tissue was then coronally sectioned using a Precisionary compresstome into 100 µm slices. Free-floating slices were incubated in Golgi impregnation solution for 7 days in the dark at room temperature. Slices were washed one week later with 0.01 M PBS-T and incubated in the remaining *sliceGolgi Kit* solutions. Striatal slices were next mounted onto gelatin-coated slides, allowed to dry completely, then dehydrated in graded ethanols, cleared using xylene and coverslipped with Permount mounting medium (Thermo Fisher Scientific, catalog #SP15-500). Golgi-stained dendrites from striatal neurons were imaged using an Axio Observer Z1 (Zeiss) microscope with a 63x/1.4NA oil-immersion objective (Zeiss). Image analysis was conducted using FIJI (Image J), where spine lengths and head widths were manually traced, and their measurements were quantified. Spine morphology was categorized using the following parameters: length less than 1 µm = stubby spine; length greater than 1 µm and width less than 0.5 µm = thin spine; length greater than 1 µm and width greater than 0.5 µm = mushroom spine (Risher et al., 2014).

### Histological Staining

Mice were perfused intracardially using 4% paraformaldehyde to achieve fixation, and whole brains were dissected. Mice were held under anathesia during the fixation procedure by the use of isoflurane (3% induction, maintained at 5% during perfusion). Fixed brains remained in 4% paraformaldehyde for one week and then briefly in 10% normal buffered formalin. Brains were then dehydrated starting with 70% ethanol, followed by 80%, 95% and absolute ethanol, followed by 100% xylene before paraffin embedding. Paraffin-embedded brain tissue was then processed into 8µm coronal sections using an automated vacuum infiltration tissue processor (TissueTek V.I.P. 5, Sakura Finetek USA). For the H&E protocol, slides were dipped in two changes of xylene for 5 minutes each. They were then re-hydrated, starting with absolute ethanol, followed by 95%, 80%, and 70% ethanol and then washed with tap water for 2 minutes. Following rehydration, slides were stained with Mayer’s hematoxylin solution (Millipore Sigma, catalog #109249) for 10-15 minutes and rinsed with tap water for 2 minutes. Slides were placed in Scott’s tap water substitute for 2 minutes and washed with tap water for 5 minutes. Next, slides were counterstained in eosin for 15 seconds to 2 minutes. Slides were then dehydrated with 95% ethanol twice for 30 seconds each, followed by 30 seconds in absolute ethanol. Slides were cleared in 3 changes of xylene, 1 minute each, and mounted with xylene-based mounting medium and coverslipped. Nuclei were stained blue, and cytoplasmic compartments were stained various shades of pink, which were used to identify tissue components. For the cresyl violet staining protocol, sections were deparaffinized and re-hydrated as described above. Slides were then stained with cresyl violet stain solution (1%) (Abcam, catalog #ab246817) for 3-5 minutes and rinsed with distilled water. Slides were then dehydrated with absolute ethanol, cleared with xylene, mounted and coverslipped as above. Nissl substance was stained violet, which was used to identify neuronal cell bodies. Images of H&E and cresyl violet stained slides were taken using an EVOS M5000 microscope (Thermo Fisher Scientific) with a 20x/0.8NA objective (Olympus). Neuropathology based on these histology stains was qualitatively assessed by a blind reviewer.

### Immunohistochemistry

A IHC hydrophobic barrier was drawn around striatal slides using a DAKO Pen (Agilent Technologies, catalog #s200230-2) which were then washed briefly in 1xPBS solution. Striatal slides were then submerged in a blocking solution consisting of 5% bovine serum albumin and 0.2% Triton-X in 1xPBS for 1 hour at room temperature. After blocking, primary antibodies (chicken anti-GFP, 1:500, Abcam, catalog #ab13970; mouse anti-GFAP, 1:500, Millipore Sigma, catalog #MAB3402; rabbit anti-IBA-1, 1:500, Cedarlane, catalog #019-19741) were diluted in a 1% bovine serum albumin/0.2% Triton-X PBS solution and incubated overnight on at 4 ℃. On the second day, slides were washed with 1xPBS and secondary antibodies (Alexa Fluor 488, goat anti-chicken, 1:500, Invitrogen, catalog #A-11039; Alexa Fluor 594, goat anti-rabbit, 1:500, Invitrogen, catalog #A-11037; Alexa Fluor 647, goat anti-mouse, 1:500, Invitrogen, catalog #A11029) diluted in a 1% bovine serum albumin/0.2% Triton-X PBS solution and incubated in the dark for 2 hours at room temperature. Slides were then washed with 1xPBS and coverslipped with DAPI Fluoroshield mounting medium to visualize nuclei (Abcam, catalog #ab104139). All images were taken using either a Axio Observer Z1 (Zeiss) microscope with a 20x/0.8NA objective (Zeiss) or a EVOS M5000 (Thermo Fisher Scientific) microscope with a 20x/0.8NA objective (Olympus).

Consistent imaging settings were used within each experiment. For GFAP and IBA-1 analysis, percent positive area values were quantified by applying a constant threshold value to all images. Percent positive area and mean gray (intensity) values were calculated using FIJI (Schindelin et al., 2012).

### Behaviour

### Open field test (OFT)

All testing was done between 8:00 A.M. and 5 P.M. In all behavioural tests described, testing times were counterbalanced among groups. All equipment was washed with 70% ethanol following each test. For OFT, mice were videotaped during a 10-minute exploration of a brightly lit open field within a 50 cm × 50 cm white Plexiglass box. Distance travelled and mean velocity were scored using ANYMaze software (Wood Dale, IL, USA). Animals with <2 cm/s mean velocity were excluded from analysis for failing to explore the arena. Trials in which ANYMaze failed to track the animal more than 10% of the time were excluded from analysis.

### Accelerated Rotarod

Mice learned to run when placed on a rotating rod (rotarod) that accelerated from 4 to 40 rotations per minute for 300 seconds. When they fell, the experimenter stopped the rotarod manually. For some trials, the mice rotated around the rotarod instead of falling, which was considered equivalent to a fall. Mice performed three trials/day for 4 days with a 1.5- to 2-hour intertrial interval. Latency to fall, number of rotations (around the rod) and number of turn arounds (where the mouse faces backwards) were recorded.

### Statistics

All statistical tests were performed using version 10.2.2 GraphPad Prism. Animal number (N) and cell/image/slice number (n) values, specific statistical tests used, p values, F values and degrees of freedom are described in Table 1. Outliers were identified using the ROUT method (Q = 1%). P values less than 0.05 were considered significant.

**Table 1.** Statistical tests used and accompanying values.

| *Figure #* | *Experiment* | *Animal (N) and cell/image/slice number (n)* | *Statistics performed* | *p value* | *Degrees of freedom/ F (DFn, DFd)* |
| --- | --- | --- | --- | --- | --- |
| 1C | HTT western blot | N=4, hemisphere n = 8 | Unpaired parametric t test | CTL vs cKO: p = 0.0061 | 1.646 (6, 5) |
| 2C | Spine density | N = 4, hemisphere n = 4, dendrite n = 26 | Unpaired parametric t test | CTL vs cKO: p = 0.4583 | (1.122) 25, 25 |
| 2C | Number of thin spines | N = 4, hemisphere n = 4, dendrite n = 26 | Mann-Whitney test | CTL vs cKO: p = 0.1859 | Not reported |
| 2C | Number of stubby spines | N = 4, hemisphere n = 4, dendrite n = 26 | Mann-Whitney test | CTL vs cKO: p = 0.6129 | Not reported |
| 2C | Number of mushroom spines | N = 4, hemisphere n = 4, dendrite n = 26 | Mann-Whitney test | CTL vs cKO: p = 0.3819 | Not reported |
| 2D | DAPI nuclear density | N=4, hemisphere n = 8 | Unpaired parametric t test | CTL vs cKO: p = 0.0865 | (3.444) 15, 14 |
| 3B | GFAP intensity | N=4, hemisphere n = 8 | Mann-Whitney test | CTL vs cKO: p = 0.0085 | Not reported |
| 3B | GFAP % area | N=4, hemisphere n = 8 | Mann-Whitney test | CTL vs cKO: p = 0.0750 | Not reported |
| 3B | IBA-1 intensity | N=4, hemisphere n = 8 | Unpaired parametric t test | CTL vs cKO: p = 0.6575 | 5.668 (15, 15) |
| 3B | IBA-1 % area | N=4, hemisphere n = 8 | Mann-Whitney test | CTL vs cKO: p = 0.2352 | Not reported |
| 4B | IV plot | N = 4, cell n = 13-22 | Two-way ANOVA Repeated measures (mixed effects) | Genotype (CTL vs cKO): p = 0.0003 | Genotype: 16.65 (1, 33) |
| 4C | Rheobase | N = 4, cell n = 13-22 | Mann-Whitney test | CTL vs cKO: p = 0.0001 | Not reported |
| 4D | Action Potential Threshold | N = 4, cell n = 13-22 | Unpaired parametric t test | CTL vs cKO: p = 0.0002 | (1.755) 12, 21 |
| 4E | Resting membrane potential (RMP) | N = 4, cell n = 13-22 | Mann-Whitney test | CTL vs cKO: p = 0.0192 | Not reported |
| 4F | Membrane resistance | N = 4, cell n = 13-22 | Mann-Whitney test | CTL vs cKO:<br>p = 0.0056 | Not reported |
| 4G | Membrane capacitance | N = 4, cell n = 13-19 | Outliers excluded,<br>unpaired parametric<br>t test (cleaned data) | CTL vs cKO:<br>p = 0.5264 | (6.374) 12,<br>18 |
| 4I | Frequency | N = 4, cell n = 12-19 | Mann-Whitney test | CTL vs cKO:<br>p = 0.1770 | Not reported |
| 4J | IEI | N = 4, cell n = 12-19 | Mann-Whitney test | CTL vs cKO:<br>p = 0.1770 | Not reported |
| 4K | Amplitude | N = 4, cell n = 12-19 | Mann-Whitney test | CTL vs cKO:<br>p = 0.0585 | Not reported |
| 5B | OFT average distance | N = 20 | Mann-Whitney test | CTL vs cKO:<br>p = 0.5117 | Not reported |
| 5B | OFT average speed | N = 20 | Mann-Whitney test | CTL vs cKO:<br>p = 0.5244 | Not reported |
| 5C | Accelerating rotarod – latency to fall | N = 20 | Two-way ANOVA | Genotype (CTL vs cKO):<br>p = 0.8883 | 0.01999 (1, 38) |

## Results

### Experimental design and validation of huntingtin conditional knockout in the striatum

To investigate the consequences of wtHTT knockout in adult striatal neurons, synapsin-promoted AAV-eGFP-Cre or AAV-eGFP was injected into the dorsal striatum of 2–4-month-old male and female *Htt*^fl/fl^ mice. All experiments took place 1-2 months post-injection and brains were assessed using histological, immunohistochemical electrophysiological and behavioural methods (Fig. 1A). Viral targeting was confirmed by IHC, where GFP was expressed widely throughout the mouse dorsal striatum, indicating the injection delivery strategy was successful (Fig. 1B). cKO of the wtHTT protein was validated by Western blot methods where Cre-injected animals had a 43.6% reduction in HTT levels on average from total brain tissue lysate 1-2 months post-surgery (Fig. 1C, p = 0.0061). A complete depletion of HTT was not expected in these total protein samples, as the tissue samples used included many non-neuronal cells that would still express normal levels of HTT. These results indicated that wtHTT levels were successfully depleted in the dorsal striatum of these animal models.

**Figure 1.**
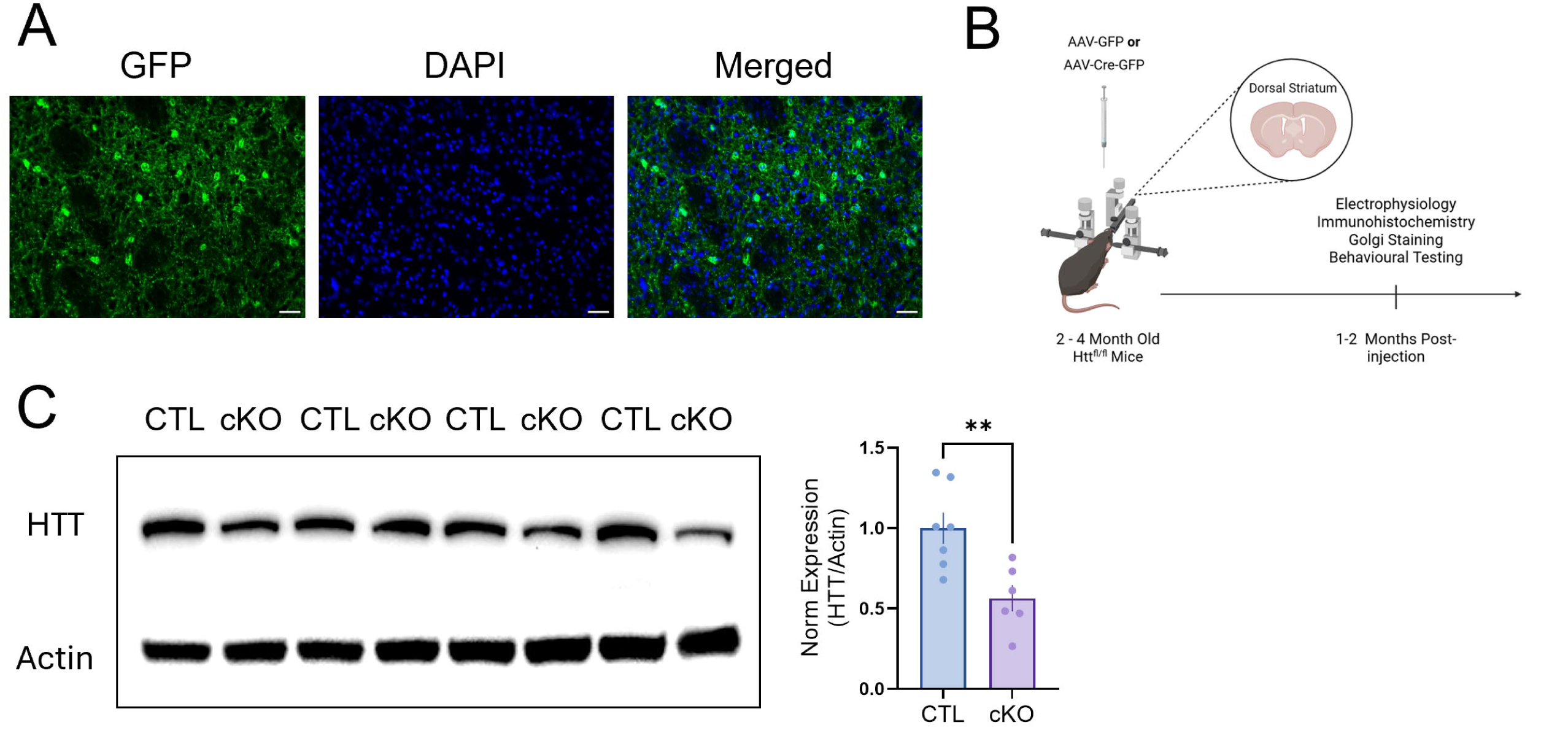
Experimental design and striatal conditional knockout mouse model validation. A Representative images showing AAV-eGFP expression and DAPI stained nuclei. Scale bar: 25 μm. B Experimental methods and timeline. C Left: Total protein samples from dorsal striatum used to assess HTT levels by Western blot. Right: Normalized HTT protein levels relative to actin control values. Data are represented as mean ± SEM.

### Huntingtin knockout does not impact general morphology in the adult striatum but leads to astrogliosis

We first wanted to determine how deletion of the wtHTT protein in adulthood would affect the general morphology of the striatum, as it was previously found that 1-2 months of wtHTT deletion in the adult mouse hippocampus had profound adverse effects on CA1 tissue organization (Barron et al., 2025). Here, control and cKO H&E-stained slides were qualitatively assessed for changes in tissue organization and cell density, as well as for signs of inflammation or neurodegeneration. No obvious signs of neuropathology could be detected with either H&E staining (Fig. 2A) or cresyl violet staining (Fig. 2B). Next, total cell density was investigated using nuclear marker DAPI to assess if wtHTT deletion impacted cell counts. These experiments showed that wtHTT cKO had only a non-significant trend towards increased nuclear density in the dorsal striatum (Fig. 2D, p = 0.0865). Spine density and morphology was also assessed by Golgi methods (Fig. 2C). Based on the Golgi experiments, no observable differences were seen between spine morphology of control and wtHTT knockout striatal neurons (Fig. 2C, spine density p = 0.458, thin spines p = 0.186, stubby spines p = 0.613, mushroom spines p = 0.382). Overall, 1-2 months wtHTT reduction did not impact tissue morphology or spine dynamics in the adult mouse brain.

**Figure 2.**
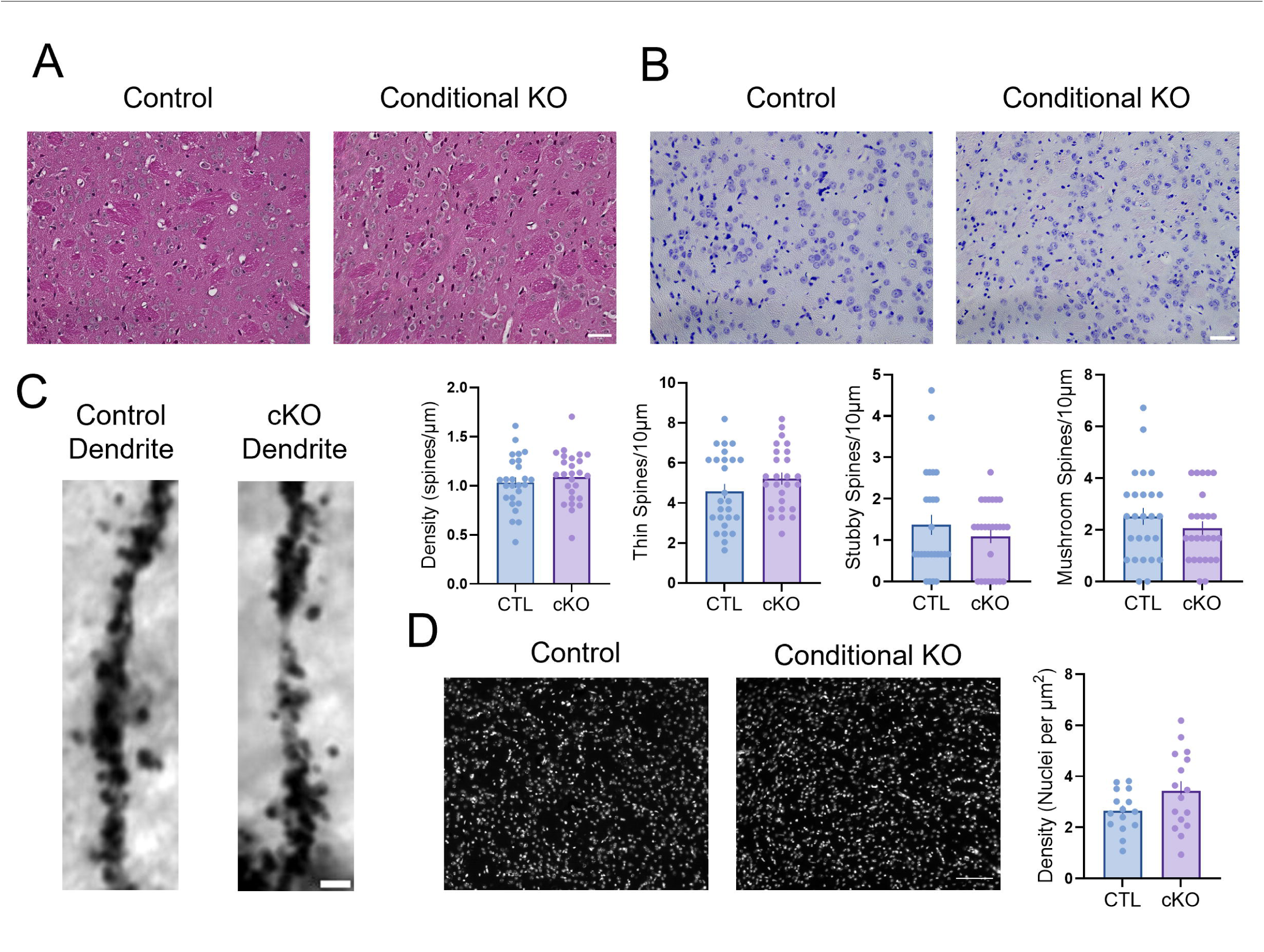
Neurohistolology of huntingtin knockout in the mouse dorsal striatum. A Representative control and wtHTT cKO slices histologically stained with hematoxylin and eosin (H&E) Scale bar: 30 μm. B Representative control and wtHTT cKO slices histologically stained with cresyl violet. Scale bar: 30 μm. C Left: Representative images of Golgi-stained dendrites from control and wtHTT cKO striatal slices. Scale bar: 2 μm. Right: Individual data and average spine density, number of thin, stubby and mushroom spines from 10 μm dendritic segments. D Left: Representative images of control and wtHTT cKO slices immunostained with DAPI. Right: Analysis of nuclear density. Scale bar: 50 μm. Data are represented as mean ± SEM.

The HD striatum is marked by significant gliosis, which correlates with disease progression (Vonsattel et al., 1985). Here, 1-2 month wtHTT cKO in striatal neurons also resulted in increased GFAP intensity (Fig. 3A; Fig 3B, p = 0.0085) and a trend towards increased GFAP percent positive area (Fig. 3D, p = 0.0750). However, quantification of IBA-1 immunostaining did not yield any significant differences in overall intensity or percent positive area values (Fig. 3C, p = 0.658; Fig. 3E, p = 0.235). Results here show that 1-2 month wtHTT knockout in the dorsal striatum leads to reactive astrogliosis, while microglia levels remain unchanged in these animals.

**Figure 3.**
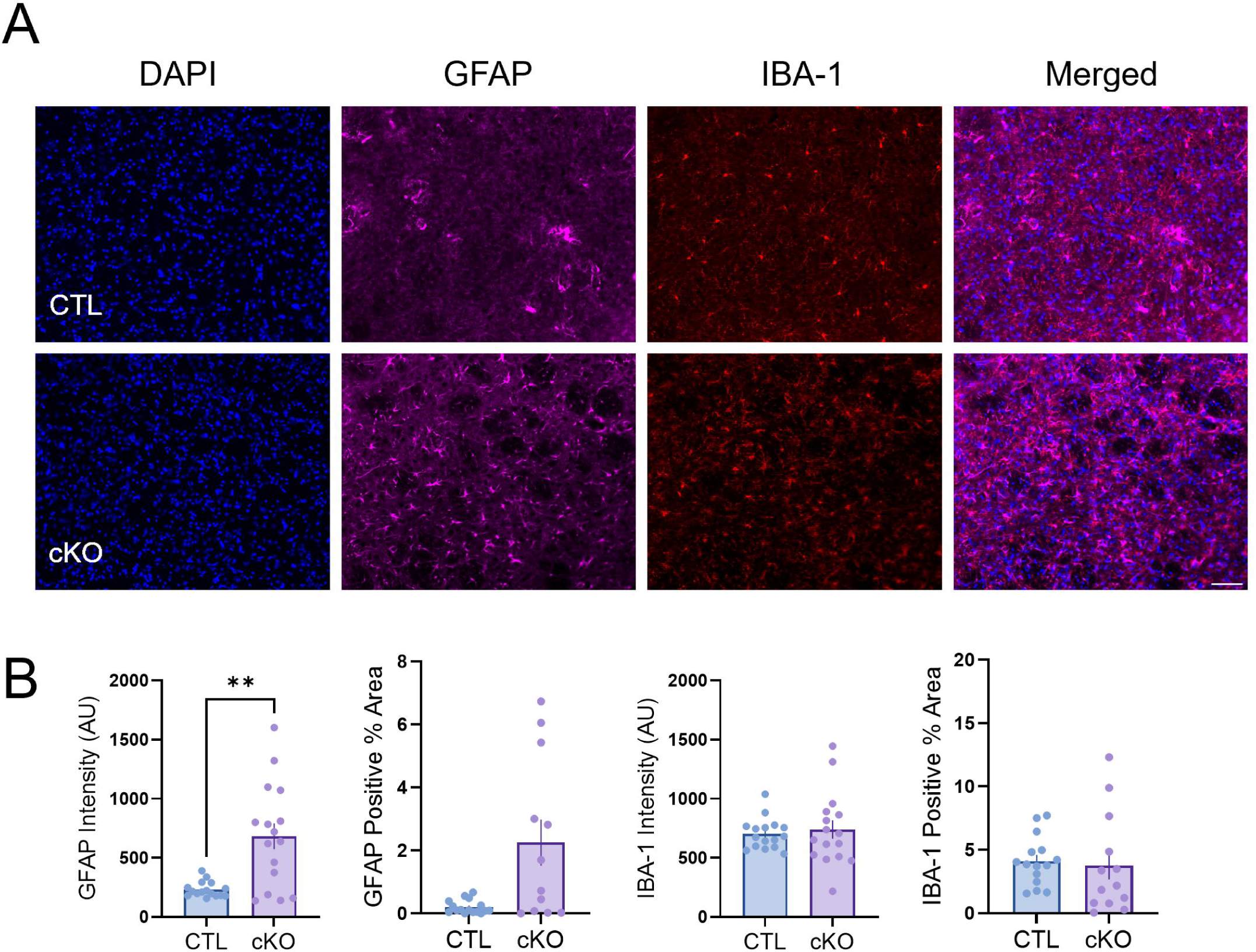
Huntingtin knockout results in astrocyte gliosis in the dorsal striatum. A Representative images of control and wtHTT cKO slices immunohistochemically stained for glia analysis. Scale bar: 50 μm. B Average GFAP staining. C Percentage of FOV with GFAP positive signal. D IBA-1 staining intensity. E Percentage of FOV with IBA-1 positive signal. Data are represented as mean ± SEM.

### Deletion of huntingtin decreases intrinsic excitability in the mouse striatum

Next, whole-cell patch clamping was used to investigate changes in cell excitability after wtHTT cKO in striatal neurons. wtHTT deletion in the striatum resulted in extensive changes in intrinsic cell properties (Fig. 4A-G). wtHTT cKO neurons fired much fewer action potentials (APs) compared to their control GFP counterparts (Fig. 4B, p = 0.0003), which was accompanied by an increased rheobase (Fig. 4C, p = 0.0001) and AP threshold (Fig. 4D p = 0.0002), indicating more positive current was required to elicit an AP. Other basic neuronal properties were impacted by wtHTT loss, including a hyperpolarized resting membrane potential (RMP) (Fig. 4E, p = 0.0192) and a decrease in membrane resistance (Fig. 4F, p = 0.0056), while membrane capacitance remained unchanged, suggesting no changes in cell size (Fig. 4G, p = 0.5264). Changes in sEPSCs in striatal neurons were also assessed after wtHTT loss. Data from these recordings show that the average frequency of spontaneous events and interevent interval (IEI) values were unchanged after wtHTT loss of function (Fig. 4I, p = 0.177; Fig. 4J, p = 0.177). However, a trending increase in sEPSC amplitude was observed (Fig. 4K, p = 0.0585). In sum, wtHTT deletion in the adult striatum resulted in profound changes in intrinsic cell excitability, while excitatory neurotransmission remained generally unaffected.

**Figure 4.**
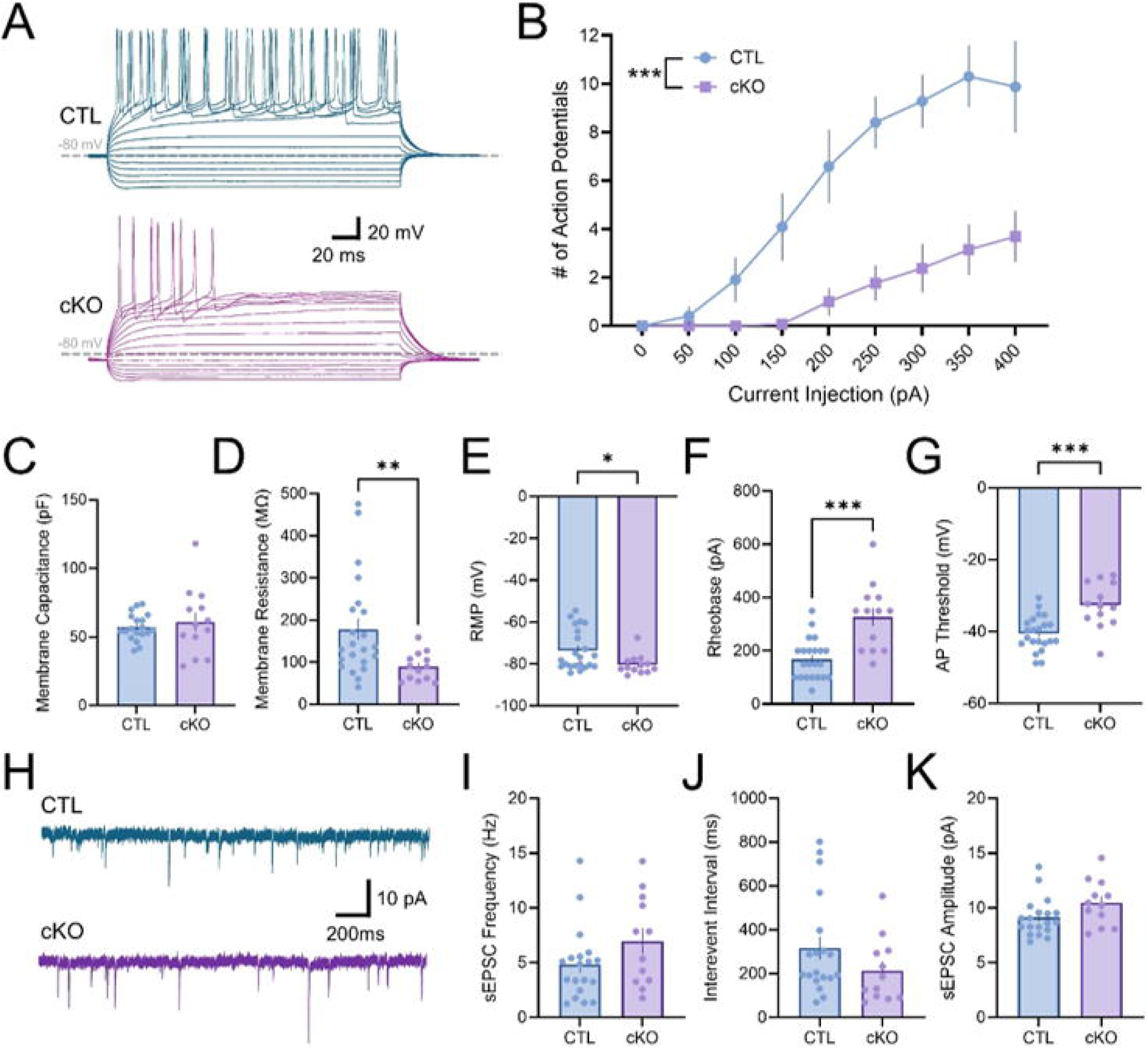
Deletion of huntingtin reduces intrinsic excitability in the mouse striatum. A Representative IV traces of control and wtHTT cKO neurons. B Current-voltage curve showing average number of APs generated at each current sweep. C Average rheobase. D Average AP threshold. E Average RMP. F Average membrane resistance. G Average membrane capacitance. H Representative sEPSC traces of control and wtHTT cKO neurons (axes: x pA, y second). I Average sEPSC frequency. J Average IEI. K Average sEPSC amplitude. Data are represented as mean ± SEM.

### 1-2 month huntingtin knockout in the mouse striatum does not influence general motor activity or motor learning

Lastly, we assessed the impact of wtHTT deletion on motor function. To do this, overall motor activity of control and cKO mice was quantified using an open field test (OFT) paradigm where mice were tracked using ANYmaze software (Fig. 5A). The same datasets but separated by sex can be viewed in supplementary data (Fig. S1). Average total distance and average speed during the ten-minute testing trial were found to be comparable between control and wtHTT cKO mice, indicating that HTT depletion in striatal neurons did not impact general motor activity in these animals (Fig. 5B, p = 0.512; Fig 5C, p = 0.5244). Next, mice were trained to run on an accelerating rotarod. This behavioural test is commonly used to quantify motor skill learning, and HD mice have significant impairments in rotarod motor learning (Van Raamsdonk et al., 2005b). For these experiments, mice were trained three times a day for 4 days, and latency to fall during each session was recorded. Both wtHTT cKO and control animals learned this behavioural test, with females performing statistically better overall compared to males (Fig. S1B, p = 0.0062). However, no differences were seen in average latency to fall over the training sessions (Fig. 5C, p = 0.888). In all, these data show that motor behaviour was comparable between control and striatal cKO animals.

**Figure 5.**
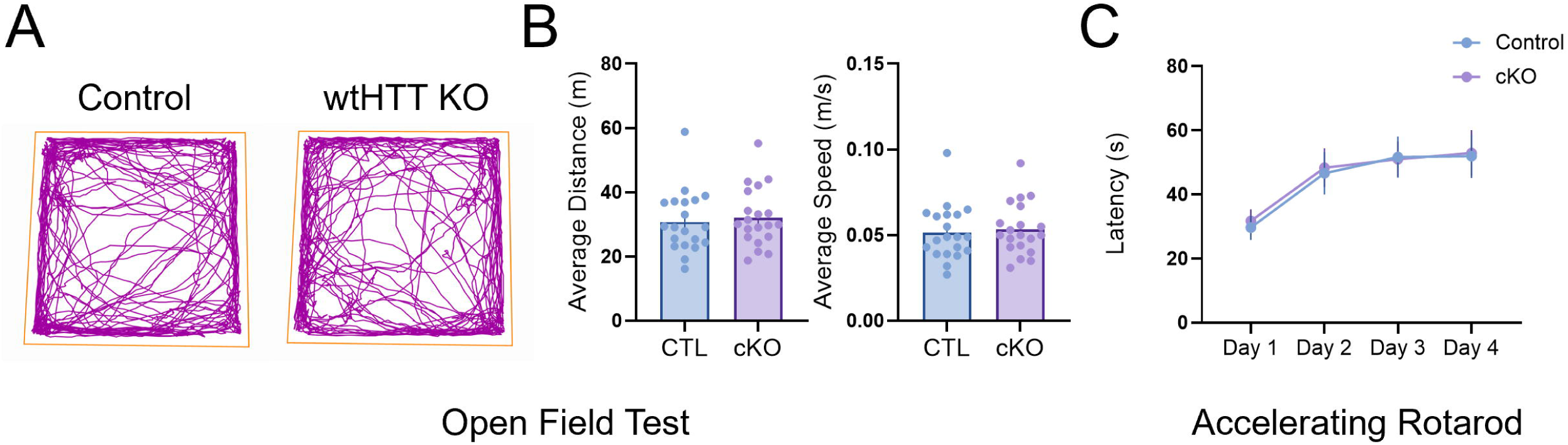
Huntingtin conditional knockout in the mouse striatum does not influence general motor activity or motor learning. OFT A Representative traces of ANYmaze video tracked movements of control and wtHTT cKO mice. B Left: Average total distance travelled. Right: Average speed. Accelerated rotarod C Average latency to fall on rotarod.

## Discussion

Results presented here demonstrate that wtHTT deletion in adult MSNs leads to reactive gliosis and decreased intrinsic neuronal excitability, while tissue morphology, spine dynamics and cell density in the striatum remain unchanged after 1-2 months wtHTT loss of function. Additionally, no differences in general motor activity or motor learning were observed after conditional wtHTT deletion in these animals. Although previous HTT-lowering studies have reported that wtHTT deletion is well tolerated in the non-human primate brain (McBride et al., 2011; Grondin et al., 2012), significant inflammatory and electrophysiological changes were discerned in these conditional knockout mouse models that may have been overlooked in previous studies evaluating the role of wtHTT in the striatum.

### Huntingtin deletion in the dorsal striatum does not impact brain morphology

Striatal neurodegeneration is a pathological hallmark of HD. The caudate and putamen structures that together make up the striatum of the basal ganglia undergo extensive volume loss, mainly due to the death of GABAergic SPNs, which comprise roughly 95% of neurons in this brain region (Vonsattel et al., 1985; G. Vonsattel and DiFiglia, 1998). Paradoxically, although HTT is ubiquitously expressed throughout the brain, striatal neurons have surprisingly low levels of HTT in health and disease states compared to other brain areas, such as the cortex and the CA2/CA3 hippocampus (Reiner et al., 2003). Striatal SPNs are especially vulnerable to the expression of mutant HTT, and many hypotheses regarding this distinct susceptibility have been reviewed elsewhere (Han et al., 2010). One prominent theory is that SPNs are discordantly impacted by transcriptional dysregulation in HD and undergo significant loss of cell type identity (Malaiya et al., 2021; Obenauer et al., 2022; Matsushima et al., 2023). Based on their unique susceptibility to cell death in HD, it was anticipated that wtHTT deletion would lead to atrophy that would be detectable at the tissue level. Surprisingly, wtHTT-deficient striatal neurons in this model showed no differences in qualitative analysis of tissue morphology and quantitative analysis of nuclear density compared to controls. Similar studies have also reported that striatal morphology is unchanged after tamoxifen-induced wtHTT-lowering in wild-type mice (Dietrich et al., 2017). Transcriptional dysregulation in the HD striatum may, therefore, be driven by a gain-of-function mechanism as opposed to wtHTT loss-of-function. Regio *et al*. recently found that when wtHTT was deleted by Cas9 gene editing in adult wild-type FVB mice, transcriptional profiles of these animals were comparable to controls (Regio et al., 2023). Striatal brain morphology was also normal in these animals, similar to what was observed here (Regio et al., 2023). We have previously shown that wtHTT reduction in primary hippocampal neurons results in a loss of transcriptional repression and altered chromatin morphology (Barron et al., unpublished observations). Still, perhaps these effects differ regionally between the hippocampus and striatum in adulthood. It will be of interest for future studies to investigate the implications of wtHTT reduction in the context of transcriptional dysregulation in the adult hippocampus.

### Reactive astrogliosis but not microgliosis in the dorsal striatum after 1-2 month huntingtin knockout

Starting early in the HD disease course, there is an increase in intensity levels of GFAP and IBA-1, astrocyte and microglia markers, respectively, and these glial cell types exhibit morphology deficits (Dowie et al., 2014). In line with this, HD patients and mouse models also display elevated levels of inflammatory cytokines and chemokines (Björkqvist et al., 2008; Wild et al., 2011). wtHTT loss of function has been previously implicated in HD neuroinflammation. For instance, wtHTT-lowered human-derived macrophages are functionally impaired and are more vulnerable to stress (O’Regan et al., 2020). Additionally, (Dietrich et al., 2017) have shown that 9 months of global wtHTT deletion in adulthood results in extensive astrogliosis. Here, 1-2 months of wtHTT cKO was sufficient to elicit widespread astrocyte reactivity in the mouse striatum. However, although it was previously found that wtHTT deletion in the hippocampus increased IBA-1 presence (Barron et al., 2025), no changes in microglial levels were detected in the current study. It is possible that the onset of microgliosis was simply missed given that microglia are one of the first immune cell types to respond to injury in the brain, with astrocytes becoming active much later (Fawcett and Asher, 1999). Astrocytes form close associations with neurons to create tripartite synapses and are important regulators of synaptic neurotransmission (Araque et al., 1999). It was recently proposed that astrocytes may also regulate intrinsic excitability of neurons, as activation of hippocampal astrocytes led to a decrease in neuronal intrinsic excitation through adenosine signalling (Expósito et al., 2024). This could be a possible mechanism underlying the change in neuronal excitability seen in both HTT conditional knockout mouse models.

### Huntingtin deletion reduces intrinsic excitability of adult striatal neurons

wtHTT cKO in the adult mouse striatum resulted in a major deficit in intrinsic excitation. Striatal SPNs receive extensive excitatory inputs from neighbouring brain areas, mainly in the form of glutamatergic signalling from the cortex and thalamus and dopaminergic signalling from the ventral tegmental area and substantia nigra pars compacta (Nelson and Kreitzer, 2014). These neurons then project through direct pathway SPNs or indirect pathway SPNs to regulate voluntary and involuntary movement, respectively. Experiments used herein did not differentiate between direct and indirect pathway SPNs. However, embryonic wtHTT deletion in either of these neuronal subsets has been previously shown to impact the inhibitory activity of globus pallidus neurons (Burrus et al., 2020). The decrease in SPN firing seen here could partly explain the change in inhibitory activity previously seen within these synaptic targets in the globus pallidus. Dorsal striatal neurons express both AMPARs and NMDARs to receive glutamatergic input from the cortex and thalamus (Jeun et al., 2009). wtHTT is known to influence AMPAR responses, likely by promoting their transport to the synapse (Mandal et al., 2011; Wennagel et al., 2022). Non-pathogenic HTT may also impact the trafficking of NMDARs through its association with HIP14 and HIP14-like proteins (Huang et al., 2011). Although sEPSC events from control and wtHTT cKO striatal neurons were statistically comparable, a trending increase in sEPSC amplitude was noted (p = 0.0585), indicating that AMPAR levels may be altered. Given that direct and indirect pathway SPNs are distinct in their postsynaptic receptor expression, downstream effects on motor activity, and pathogenesis in HD, it is possible that changes in spontaneous excitatory activity differed between these striatal cell types. Therefore, the present study may have been unable to detect these cell type-specific changes (Kravitz et al., 2010; Oldenburg and Sabatini, 2015; Lee et al., 2016; Barry et al., 2018). The impact of wtHTT loss in adulthood on striatal excitation remains an important area of investigation, given the critical role of the striatum in basal ganglia synaptic circuitry and HD. Although changes in motor activity were not observed in these experiments, previous studies investigating this have yielded mixed results. Global deletion of wtHTT between 3-9 months in mice elicited progressive motor deficits, including decreased latency to fall from an accelerated rotarod (Dietrich et al., 2017). While wtHTT knockout more specifically targeted to the adult striatum and projecting areas showed no behavioural deficits in a comprehensive motor assessment (Regio et al., 2023). Therefore, the previously identified effects on motor performance may be due to wtHTT-lowering in another brain area downstream from the striatum, such as the cortex.

Overall, wtHTT conditional deletion in adult striatal SPNs decreased intrinsic neuronal excitability and produced an astrogliosis response, while tissue organization, spine morphology, cell density and motor phenotypes remained unaffected. Results presented here contribute additional evidence to the growing body of literature that demonstrate wtHTT loss of function negatively impacts neuronal health in the adult brain and caution that substantial reduction of wtHTT levels through the use of non-selective HTT-lowering therapeutics may have adverse molecular consequences such as those identified here.

## Acknowledgements

We would like to thank Dr. Scott Zeitlin (University of Virginia) for providing us with our initial Htt^fl/fl^ mouse colonies that were bred and used for all experiments. We would also like to thank the Animal Care Services staff at Memorial University for their assistance with the housing, monitoring and care of all research animals used for these experiments. All research presented in the current manuscript was funded by Dr. Parsons’ Canadian Institutes of Health Research (CIHR) project grant.

## Author Contributions

JCB and MPP designed research; JCB, MLG, EPH, FA, FN, CSM and MPP performed research; JCB analyzed data; JCB wrote the paper with feedback from MPP.

---

**Figure S1.**
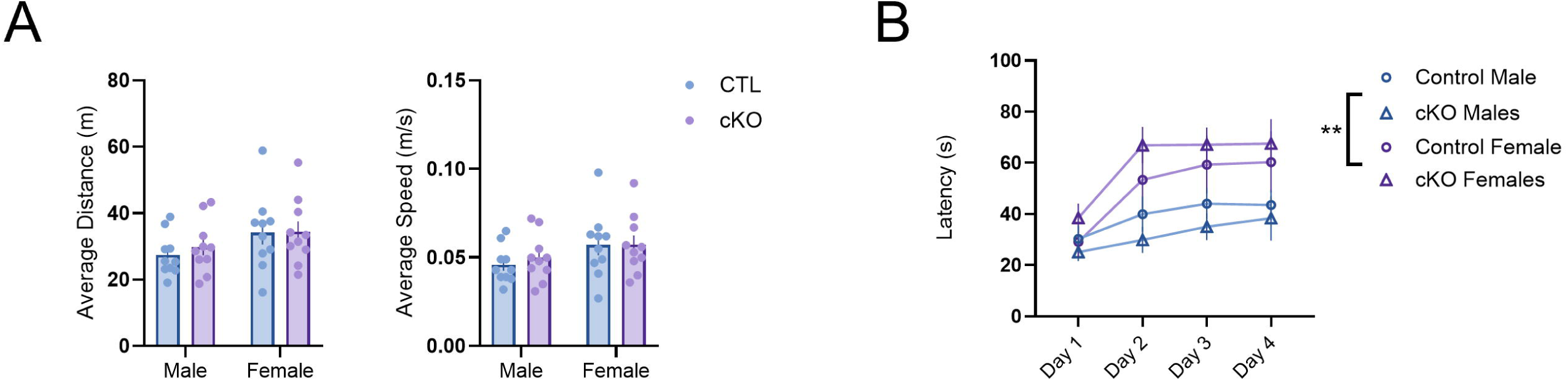
A OFT data separated by sex. B Rotarod data separated by sex. Data were assessed for sex differences by two-way repeated measures ANOVA (B) and two-way ANOVA and are represented as mean ± SEM.

